# Beyond temperature: Relative humidity systematically shifts the temperature dependence of population growth in a malaria vector

**DOI:** 10.1101/2025.05.30.656372

**Authors:** Paul J. Huxley, Joel J. Brown, Brandy St. Laurent, Britny Johnson, Olivia Y. Cheung, Anna Asamoah, Brandon D Hollingsworth, Eric R. Bump, Michael C. Wimberly, Mercedes Pascual, Leah R. Johnson, Courtney C. Murdock

## Abstract

Understanding ectotherm responses to environmental change is central to coping with many of humanity’s current and future challenges in public health, biodiversity conservation, and food security. Complex relationships between abiotic and biotic factors can influence ectotherm abundance and distribution patterns by introducing stage–specific variation in fitness trait responses. Variation in temperature, rainfall, competition, and habitat have all been considered in previous attempts to understand how environmental factors can interact and vary in their relative influence on species’ maximal population growth rates, *r*_m_. However, the combined effects of temperature and humidity on this fundamental metric are poorly understood. We show that variation in relative humidity can influence juvenile trait responses and *r*_m_’s temperature dependence in *Anopheles stephensi*, an important malaria vector. Our climate suitability maps show that the interactive effects of temperature × humidity on juvenile traits have important implications for predicting how environmental change will influence arthropod–mediated systems.

## Introduction

Arthropods, including many disease vectors like mosquitoes, experience a complex suite of environmental factors, both abiotic (e.g., temperature, rainfall, humidity, salinity) and biotic (e.g., biological enemies, inter– and intra–specific resource competition, and variation in habitat quality). Such factors can interact and vary in their relative influence on a population’s capacity for growth, and, therefore, its abundance and distribution patterns (Huxley et al., 2022, 2021; Kleynhans and Terblanche, 2011; Evans et al., 2019; Ryan et al., 2023, 2015; Murdock et al., 2017). To better predict how environmental variation constrains current and future abundance and distribution patterns in organisms of interest, it is critical to consider the effects of multiple environmental factors and their interactions on arthropod fitness and population dynamics.

In vector ecology, as well as the broader field of ecology, there has been a strong emphasis on studying the effects of temperature due to climate change (reviewed in Mordecai et al., 2019). Body temperature has important effects on the rates of enzymatic processes as well as the structural integrity of cellular membranes and proteins (Angilletta, 2009). The effects of environmental temperature on ectotherm performance, including mosquitoes, are typically non-linear, with performance of a given rate process increasing from zero at a minimum temperature (*T*_min_) up to an intermediate temperature where performance peaks (*T*_pk_), followed by a steep decline towards a critical upper temperature where the organism typically experiences death (*T*_max_). The *T*_min_ and *T*_max_ represent the operational limits for trait performance because temperatures that exceed this range are not permissive for ectotherm survival, reproduction, or development (Corkrey et al., 2016; Deutsch et al., 2008; Hoffmann et al., 2013; Sinclair et al., 2016). These thermal limits are consistent with the Metabolic Theory of Ecology in hypothesizing that metabolic rate increases exponentially with increases in environmental temperature, and, as a consequence, the maximal population growth rate, *r*_m_, and its underlying life history traits also exhibit a similar temperature dependency (Pawar et al., 2024; Amarasekare and Savage, 2012). Collectively this information gives us a Thermal Performance Curve (TPC), which has been used widely to infer ecological and evolutionary outcomes in response to current and future temperature variation (Huey and Kingsolver, 2019; Pawar et al., 2024; Mordecai et al., 2019; Miazgowicz et al., 2020; Tesla et al., 2018). In addition to environmental temperature, water availability is another critical abiotic variable that influences ectotherm biology, and both play important roles in determining the abundance and distribution of organisms (reviewed in Brown et al., 2023; Rozen-Rechels et al., 2019). For mosquitoes that transmit human pathogens, water in the environment has multiple effects on population fitness and dynamics. Water availability is an important determinant of the carrying capacity of a given population, as juvenile mosquitoes develop in aquatic environments. For mosquitoes that live in close association with human populations and that transmit pathogens like malaria, arboviruses (e.g., dengue, chikungunya, and Zika), and filariasis, water in the environment is largely supplied by a mixture of rainfall (Abiodun et al., 2016; Althouse et al., 2015; Parham et al., 2012; Bomblies, 2012; Paaijmans et al., 2007), irrigation for agriculture (Norris, 2004; Ijumba et al., 2002) and water storage practices (Evans et al., 2019; Surendran et al., 2019; Brown et al., 2014; Stewart Ibarra et al., 2013; Padmanabha et al., 2010). For mosquitoes that develop in artificial containers (e.g., *Aedes aegypti* and *Ae. albopictus*, *Anopheles stephensi*, multiple *Culex spp*.), water storage practices and variation in humidity may be more important determinants of water availability than precipitation (Santos-Vega et al., 2016, 2022). Further, due to the fundamental relationship that exists between temperature and the amount of moisture the air can hold (Brown et al., 2023), variation in relative humidity and temperature alters evaporation rates in aquatic environments where juvenile mosquitoes develop, (Juliano and Stoffregen, 1994) as well as potentially altering intrinsic features of these environments, such as the concentration of solutes (Juliano and Stoffregen, 1994) and surface tension (Pérez-Díaz et al., 2012; Singh et al., 1957).

In this study, we investigate the influence of variation in relative humidity during juvenile development on the temperature dependence of *r*_m_ in *Anopheles stephensi*, the South Asian urban malaria vector that has recently become an important threat to malaria control efforts in Africa (Zhou et al., 2024). We conducted a large, high resolution environmental experiment where *An. stephensi* larvae were exposed to nine different constant temperature conditions (14*^◦^*C–42*^◦^*C), five constant relative humidity conditions (30%–90%), and two different evaporation scenarios. Specifically, we measured how variation in relative humidity affected the temperature dependence of juvenile trait performance (development rate, survival probability, and body size upon emergence) and, using an analytic *r*_m_ model, we calculated *r*_m_ to predict how these effects can, in turn, influence the temperature dependence of maximal population growth rate. Our study has important implications for the inferences gained from previous ecological studies that focus on the effects of temperature alone, as well as for current and future predictions of this invasive species to establish and thrive in urban centers in Africa, contributing to urban malaria transmission on the continent.

## Methods

### Colony maintenance

Establishing field derived colonies of *An. stephensi* is incredibly difficult due to the nature of its life cycle, and government restrictions in the native range limit our ability to obtain biological material on this species. Thus, here we used an *An. stephensi* urban–type strain acquired from a longstanding (∼40 years) colony at Walter Reed Army Institute of Research, via the University of Georgia. Prior to the experiments reported here, the colony was maintained using established methods described in Miazgowicz et al. (2020) and Pathak et al. (2019).

### Experimental set–up

For the experiments, we used Percival 36VL incubators with ultrasonic humidification systems and 12:12 light:dark cycles. We exposed immature mosquitoes to eight constant temperatures (16*^◦^*C, 20*^◦^*C, 24*^◦^*C, 28*^◦^*C, 32*^◦^*C, 36*^◦^*C, 38*^◦^*C, 40*^◦^*C) across five relative humidity levels (30%, 45%, 60%, 75%, and 90%), which generated 40 temperature–humidity combinations. This full–factor design was later supplemented with treatments at 14*^◦^*C and 42*^◦^*C to better capture the minimum and maximum temperature limits for trait performance, respectively. Experimental blocks contained a treatment at each temperature (Table 1). We also included an uncontrolled evaporation treatment to determine the extent to which fitness trait responses differed when competition for space was intensified as rearing environments diminished in size. Larvae were held within the incubators in three replicate plastic trays (Sterilite 5.7 litre; 35cm x 21cm x 12cm) for each treatment (controlled and uncontrolled evaporation) covered with fine mesh. The trays in each experimental block initially comprised of 100 first instar (L1) larvae, four pellets of Cichlid Gold fish food and 1 litre of purified water (reverse osmosis) that had been pre–warmed/cooled to the temperature of the particular treatment (i.e., a tray of larvae placed into an incubator at 35*^◦^*C was provided with water warmed to 35*^◦^*C). For the controlled evaporation trays, we recorded evaporated water volumes and replaced the water that was lost at the required temperature daily. Trays were rotated within the incubator to equalize any uneven evaporation effects. Upon pupation, pupae were removed and placed into a 250ml plastic cups containing treatment–temperature water. Pupae cups were then placed into 17.5cm × 17.5cm × 17.5cm fine mesh cages (BugDorm Small) and put into the relevant incubator. This experimental set–up generated life history trait data for 24,000 individual juvenile mosquitoes.

**Table 1:**
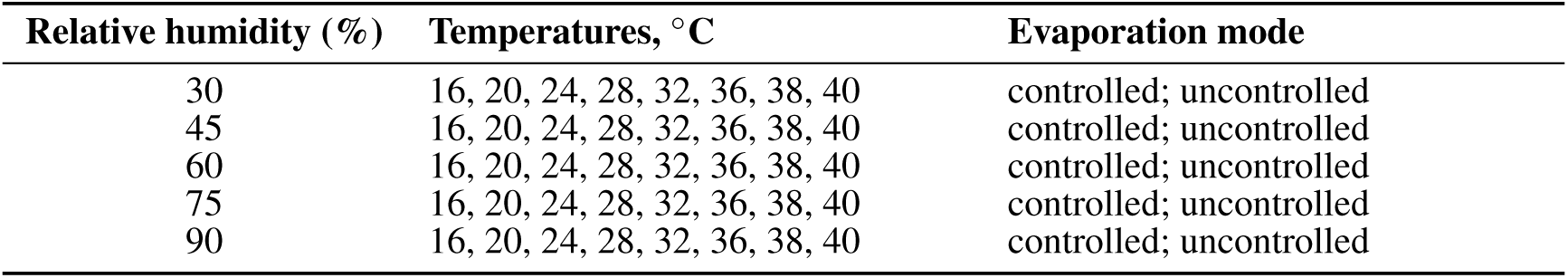
Temperature and relative humidity treatments for each experimental block.

| Relative humidity (%) | Temperatures, °C | Evaporation mode |
| --- | --- | --- |
| 30 | 16, 20, 24, 28, 32, 36, 38, 40 | controlled; uncontrolled |
| 45 | 16, 20, 24, 28, 32, 36, 38, 40 | controlled; uncontrolled |
| 60 | 16, 20, 24, 28, 32, 36, 38, 40 | controlled; uncontrolled |
| 75 | 16, 20, 24, 28, 32, 36, 38, 40 | controlled; uncontrolled |
| 90 | 16, 20, 24, 28, 32, 36, 38, 40 | controlled; uncontrolled |

### Measuring life history trait performance

During the experiment, the number of individuals emerging each day in each treatment and tray was recorded until all individuals had either emerged or died prior to emergence. From these observations, two juvenile traits were calculated. First, juvenile survival proportion was calculated for each treatment group and tray by subtracting the number of adults that emerged from each tray from the initial number of larvae placed in each tray at the start of the experiment (*n*=100). That is, any juveniles that did not emerge were assumed to have died. Second, the juvenile development time was calculated based on the number of days required for surviving individuals to emerge as adults.

Additionally, wings (one per individual) were measured in a subset of adult females upon emergence (*n*=20 per treatment per tray). Wing length measurements (mm) were taken from the tip of wing (excluding fringe) to the distal end of the alula using a dissecting scope and micrometer (Evans et al., 2018; Murdock et al., 2017, 2012).

### Calculating maximal population growth rate, ***r*_m_**

To determine how variation in relative humidity can affect the temperature dependence of mosquito population growth rate (*r*_m_), we used a continuous–time, stage–structured model based on the Euler–Lotka equation (Charnov, 1993; Savage et al., 2004; Amarasekare and Savage, 2012). Cator et al. (2020) derived an approximation appropriate for the range of growth rates typically seen across arthropods (Eqn. 1; Table 2):

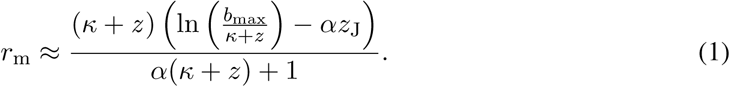

**Table 2:** Definitions of model parameters.

| Parameter | Units | Description |
| --- | --- | --- |
| $r_m$ | $\text{day}^{-1}$ | Maximal population growth rate |
| $\alpha$ | days | Hatching-to-adult development time |
| $b_{\max}$ | $\text{eggs} \times (\text{female} \times \text{day})^{-1}$ | Maximum fecundity rate |
| $\kappa$ | $\text{day}^{-1}$ | Fecundity loss rate |
| $z$ | $\text{day}^{-1}$ | Adult mortality rate |
| $p_{\text{EA}}$ | probability | Survival averaged across juvenile stages |
| $T_{\text{pk}}$ | $^{\circ}\text{C}$ or $\text{K}$ | Temperature at which trait performance peaks |
| $T_{\min}$ | $^{\circ}\text{C}$ or $\text{K}$ | Lower temperature at which trait performance ceases |
| $T_{\max}$ | $^{\circ}\text{C}$ or $\text{K}$ | Upper temperature at which trait performance ceases |
| $B_{\text{pk}}$ | Measurement unit of trait | Trait performance achieved at $T_{\text{pk}}$ |
| $T_{\text{opt}}$ | $^{\circ}\text{C}$ | Temperature at which $r_m$ peaks |
| $r_{\text{opt}}$ | $\text{day}^{-1}$ | $r_m$ achieved at $T_{\text{opt}}$ |

Here, *α* is egg–to–adult development time (days), *b*_max_ is peak reproductive rate (individuals (eggs) × individuals (females) × day^−1^), *κ* is the fecundity loss schedule (individual^−1^ day^−1^), and *z*_J_ and *z* are juvenile and adult mortality rates (individual^−1^ day^−1^), respectively. Although Eqn. 1 is an approximation, it is sufficiently accurate providing *r*_m_ is less than 1 (in units of day^−1^; Cator et al., 2020), which is typically true for insect growth rates (Frazier et al., 2006; Pawar et al., 2024). Eqn. 1 explicitly incorporates the traits underlying *r*_m_, so it can be used to analytically understand how variation in these traits can affect *r*_m_. Further details of the derivation and underlying assumptions of *r*_m_ can be found in (Cator et al., 2020).

Juvenile mortality rate (*z*_J_ in Eqn. 1) is not directly obtainable from the data collected in this study. Instead, we measure the probability of emergence as an adult (also called the egg to adult survival probability), which we notate as *p*_EA_. In Eqn. 1, the term *αz*_J_ is approximately the survival probability if the development time is sufficiently small. More specifically, this term is based on a first order Taylor’s expansion:

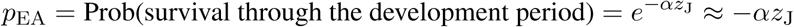

We use the original exponential relationship, *p*_EA_ = *e^−αz^*^J^, so that ln(*p*_EA_) = −*αz*_J_. Replacing −*αz*_J_ in Eq. 1 with ln(*p*_EA_) gives our target equation:

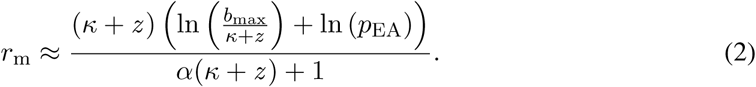

### Modelling juvenile trait thermal performance in the context of relative humidity variation

The thermal dependence of each of the five life–history traits (Table 2) in Eqn. 2 can be described by a Thermal Performance Curve (TPC). By substituting fitted TPCs into this equation, it then gives the temperature dependence of *r*_m_.

We modelled the development time TPCs for each humidity level in two ways. First, to get the juvenile development time directly, *α* in Eqn. 2 we used an exponential decay function (Supplementary Eqn. 1). This is a parsimonious approach that is consistent with the observed development time data. However, this model is not unimodal and does not have defined values for *T*_min_, *T*_pk_, *B*_pk_, and *T*_max_. Thus in order to be able to make a more direct comparison to previous studies, we fitted the standard Briere model (Supplementary Eqn. 2; as implemented in the `bayesTPC` package in R; Briere et al., 1999; Sorek et al., 2025) to *inverted development times* (i.e., development rate; 1/*α* in Eqn. 2). The temperatures at which trait performance peaks and its value at its peak (*T*_pk_ and *B*_pk_, respectively; Table 2) were estimated numerically from the posterior distributions for each humidity level’s TPC and summarized by the posterior medians and Highest Posterior Density (HPD) intervals.

The probability of egg to adult survival, *p*_EA_ described above was estimated from the juvenile survival data (number emerged out of initial number of eggs) using a binomial generalized linear model with a log link function (Supplementary Eqn. 3), as implemented in `bayesTPC` (Sorek et al., 2025). Similarly to development, to obtain all quantities of interest (*T*_pk_, *B*_pk_, *T*_min_ and *T*_max_) for juvenile survival, we must calculate these from the posterior distribution of parameters. We obtained estimates for *T*_pk_ and *B*_pk_ using the same procedure as for development rate. However, the calculated survival function asymptotes to zero (due to the link), and so there is not a true *T*_min_ and *T*_max_. Instead, we define *T*_min_ and *T*_max_ at each humidity level to be the temperatures at which survival probability was less than or equal to 1%.

Note that in this study we do not measure how variation in humidity can influence the temperature dependence of adult life history traits that contribute to *r*_m_. Thus, in our *r*_m_ calculations we held adult mortality and fecundity rates (*z* and *b*_max_, respectively in Eqns. 1, 2) constant across temperature × humidity levels at 0.12 day*^−^*^1^ and 11.37 day*^−^*^1^, respectively. These averages were estimated using a global dataset that we synthesized from the literature on the temperature dependence of these traits in *An. stephensi* (Supplementary Data file). As *κ* has been shown to only make a very small contribution to *r*_m_ (Cator et al., 2020), we assume that *b*_max_ declines with age at a constant rate of 0.01 individual^−1^ day^−1^. Finally, to obtain all quantities of interest for *r*_m_ (*r*_opt_, *T*_opt_, *T*_min_ and *T*_max_), we used a similar procedure to the one that we used for survival. However, for *r*_m_ *T*_min_ and *T*_max_, we used the TPC posteriors at each humidity level to estimate the temperatures at which *r*_m_ was zero.

### Modelling the thermal dependence of adult wing length in the context of relative humidity variation

Wing length was measured as a proxy for body size, which is itself a proxy measure of reproductive potential. Both relationships are expected to be linear—larger females have longer wings, greater body mass, and are more fecund (Briegel, 1990a,b). To analyse how relative air humidity can affect the temperature–size rule (i.e., body size generally decreases with temperature in ectotherms; Atkinson, 1995), we fitted a Bayesian hierarchical regression model to adult female wing lengths using the `brms` package in R (R Core Team, 2023; Bürkner, 2017). In this model (wing length ∼ temperature × humidity + (1 | tray)), wing length is the outcome variable and temperature and relative humidity (RH) are the predictor variables. The interaction term (temperature × humidity) tests both the individual effects of the predictor variables on wing length, as well as their combined interaction effect. The random effect variable (1 | tray) allows each tray to have its own baseline wing length, accounting for tray-specific differences, but the effect of temperature and RH on wing length is assumed to be consistent across trays. The model assumes a Gaussian distribution and it ran for 5000 iterations. The Hamiltonian Monte Carlo sampling algorithm was set to a maximum tree depth of 20.

### Trait sensitivity analysis

To determine the extent to which variation in humidity can affect the *relative* contributions of the juvenile fitness traits to *r*_m_’s temperature dependence we conducted a sensitivity analysis. This analysis uses the derivatives of *r*_m_ with respect to the traits of interest to determine the rate at which *r*_m_ changes with temperature (similarly to Mordecai et al., 2013; Cator et al., 2020). Full details of the approach are described in the Supplementary Information. Briefly, we derive an equation via the chains rule such that each summed term in the equation (Eqn. SE4) quantifies the relative contribution of each temperature dependent trait (described by the fitted TPC) in Eqn. 2 to the temperature dependence of *r*_m_ (i.e., to the overall derivative with respect to temperature). For simplicity and clarity in this calculation, we set the parameters in each TPC to the Maximum *A Posteriori* (MAP) estimator for each trait–humidity combination (Figs. 4 & S8) instead of using all posterior samples. The sample–based MAP estimator is calculated as part of the MCMC fitting process in the `bayesTPC` package (Sorek et al., 2025) in R.

### Mapping climatic suitability for mosquito population growth

To show the potential for variation in relative humidity to affect climatic suitability for *An. stephensi*, we used historical (1970–2000) gridded temperature and humidity from the NASA–NEX–GDDP– CMIP6 dataset to map climatic suitability for mosquito population growth using *r*_m_.

Daily data were first aggregated into monthly climate summaries were then used to compute monthly *r*_m_ values. These values were then averaged into quarterly means (January–March, April– June, July–September, October–December) for both South Asia and Africa. For Africa, these quarters capture shifts in the Inter-Tropical Convergence Zone (ITCZ) from its southernmost location in January–March to its northernmost location in July–September. For South Asia, these quarters align with the winter dry season (January–March), pre–monsoon (April–June), monsoon (July– September), and post-monsoon (October–December). Maps were generated for South Asia and Africa using two models: a temperature–only model that assumed 75% RH at all locations and a temperature × humidity model that allowed temperature–trait relationships to vary with humidity. The two models were differenced for each quarter to highlight geographic areas where humidity substantially affected mosquito population growth. For each model, we also calculated the total land area suitable for mosquito growth, which was defined as *r*_m_ > 0 throughout all months of the year. Further details of our mapping approach are in the Supplementary Materials.

## Results

### Relative humidity effects on temperature–dependent juvenile traits

The realised effect of temperature on juvenile trait responses was systematically mediated by relative humidity. Non–overlapping 95% HPD intervals indicate that the differences between the TPC parameters for juvenile survival probability at most humidity levels differed significantly (similar to a statistically significant difference at the *p <* 0.05 level in a classical analysis; Fig. 1a–c; Tables S1, 2). At its peak, the probability of egg to adult survival was around 11% lower at 45% RH than at 90% RH, and the temperature at which it peaked (*T*_pk_) was ∼4.5*^◦^*C lower at 30% RH than at 90% RH (Fig. 1b). At these humidity extremes, *T*_min_ was about 4*^◦^*C lower at 30% RH than at 90% RH and *T*_max_ was ∼5*^◦^*C higher at 30% RH than at 90% RH (Fig.1c; Table S2). Differences among the TPC parameters across humidity levels for juvenile development rate were also common (Fig. 1d– f; S3 & 4). At its peak, development rate was at its highest at 60% RH (0.15 day^−1^; ∼6.67 days) and it was lowest at the lower and upper RH level extremes (30% RH: 0.132 day^−1^ (∼7.58 days); 90% RH: 0.141 day^−1^ (∼7.09 days)). The development rate *T*_pk_ was ∼1.2*^◦^*C higher at 30% RH than at 90% RH (Fig. 1e; Table S3). For this trait, the greatest differences in *T*_min_ were predicted at the humidity extremes. At 30% RH, *T*_min_ was ∼8.6*^◦^*C whereas it was ∼5 degrees higher at 90% RH (∼13.6*^◦^*C). At all humidity levels above 60% RH, development rate’s *T*_max_ was ∼44.7*^◦^*C and it was ∼43.9*^◦^*C below 60% RH (Fig. 1f; Table S4).

**Figure 1:**
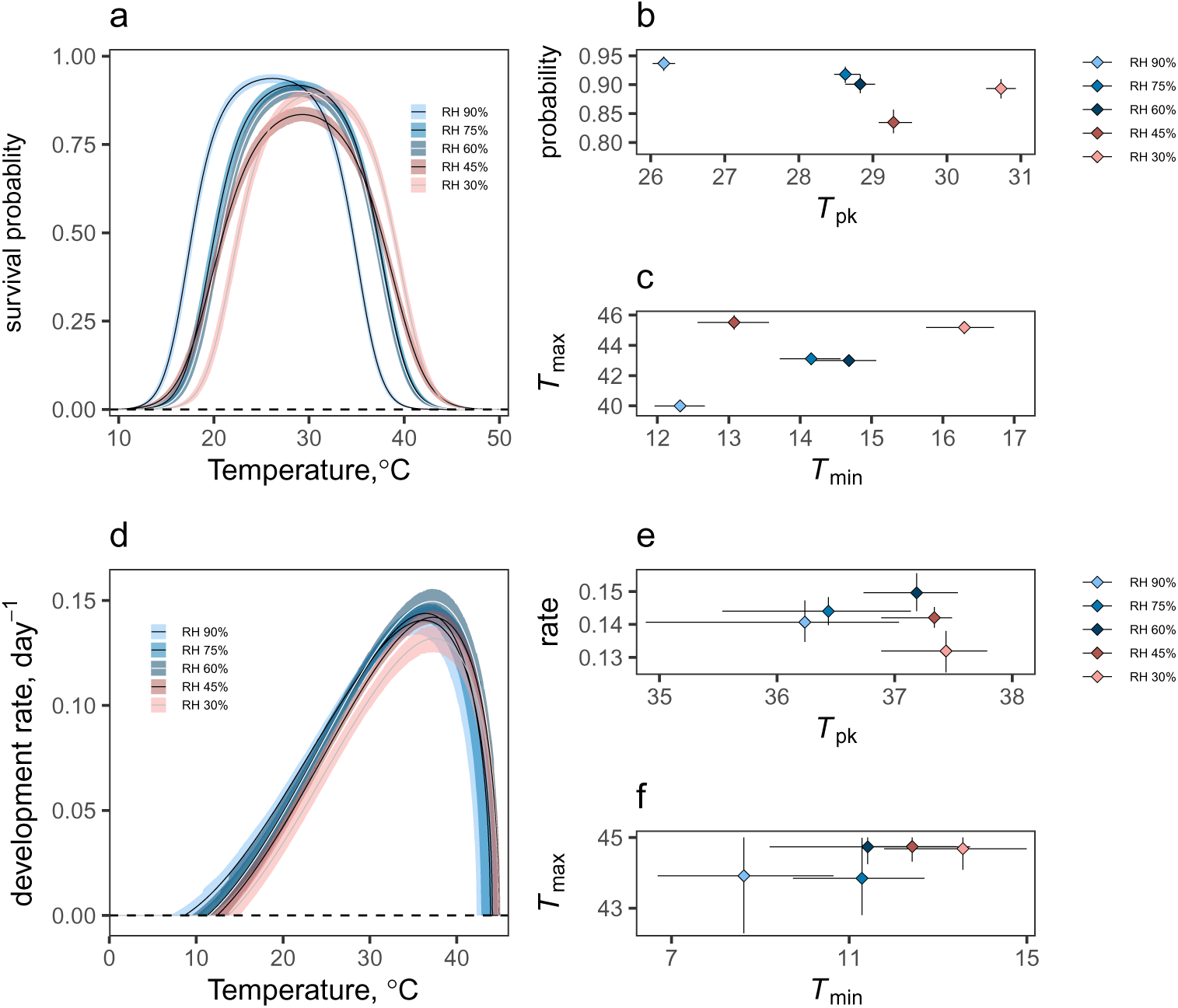
Relative humidity shapes the temperature dependence of juvenile fitness traits in *Anopheles stephensi*. The effects of relative humidity variation on the thermal performance curves (TPCs) of juvenile survival probability (*p*_EA_ in Eqn. 2), and juvenile development rate (1/*α* in Eqn. 2). Shading around the lines are HPD intervals calculated using the posteriors for each humidity–dependent TPC. Ball and stick diagrams represent how the predicted thermal peaks (*T*_pk_), and minimum (*T*_min_) and maximum (*T*_max_) vary at each relative humidity level for each trait. Bidirectional error bars are 95% CIs. **Survival Probability**, *p*_EA_: **(a)** posterior distribution of fitted TPCs across humidity levels; **(b)** maximum survival probability vs. *T*_pk_; **(c)** *T*_min_ and *T*_max_. **Development rate**, 1*/α*: **(d)** posterior distribution of fitted TPCs across humidity levels; **(e)** maximum development rate vs. *T*_pk_; **(f)** *T*_pk_ vs. *T*_max_. The development time data for observed individuals that we inverted for the development rate curves are shown in Fig. S2. In all panels, non–overlapping CIs indicate statistical significance similar to at least the *p <* 0.05 level in a classical analysis.

For the uncontrolled evaporation treatments, juveniles developing in trays held at 30% or 45% RH did not survive to adulthood regardless of temperature. However, the juvenile survival TPCs at RH levels above 60% (mortality was complete at levels below 60% RH) were similar to those observed for these RH levels in the controlled experiment (Fig. S3a–c; Tables S7 & S8). At 90% RH, survival probability was 0.95 at its *T*_pk_ (∼26*^◦^*C); it was about 6% lower at 75% RH (0.89) and 35% lower at 60% RH (0.62) than it was at the *T*_pk_ for 90% RH (∼31*^◦^*C and ∼33*^◦^*C, respectively). *T*_min_ increased from ∼11*^◦^*C at 90% RH to ∼26*^◦^*C at 60% RH; *T*_max_ was ∼40*^◦^*C at these RH levels and it was ∼42*^◦^*C at 75% RH. Without substantial prior information, it was only possible to fit development rate TPCs at the highest RH levels (75% and 90% RH) in the uncontrolled evaporation treatments (Fig. S3d–f; Tables S9 & S10). At these RH levels, development rate was ∼0.15 day^−1^ at 31.98*^◦^*C and 33.92*^◦^*C, respectively. The *T*_min_ for this trait increased from ∼13.29*^◦^*C at 90% RH to ∼14.60*^◦^*C at 75% RH; *T*_max_ was ∼37.86*^◦^*C at 90% RH and ∼39.85*^◦^*C at 75% RH. The interaction between temperature and humidity caused significant variation in adult body size in the uncontrolled treatments. Size decreased both at warmer temperatures and at lower humidity levels, but in contrast with the controlled treatments the decrease with temperature was greater in the lower humidity levels (75 and 60% RH) than at the highest humidity level (90%). As temperatures increased from 16 to 40*^◦^*C, size was predicted to decrease by ∼1.43mm at 60% RH, whereas size at 90% RH decreased by ∼0.92mm (Fig. S5).

### Relative humidity effects on temperature–dependent wing length

The temperature × humidity interaction also resulted in significant variation in size at maturity (wing length) between humidity levels (Fig. 2). Adult size decreased with temperature (Slope: –0.0216; Credible Interval: –0.0276, –0.0158), increased with humidity (Slope: 0.0063; CI: – 0.0034, 0.0092), but decreased more steeply with temperature at higher humidity levels (Interaction: –0.0002; CI: –0.0003, –0.0001). As temperatures increased from 16 to 40*^◦^*C, size was predicted to decrease by ∼1mm at 90% RH, whereas at 30% RH it was predicted to decrease by ∼0.07mm.

**Figure 2:**
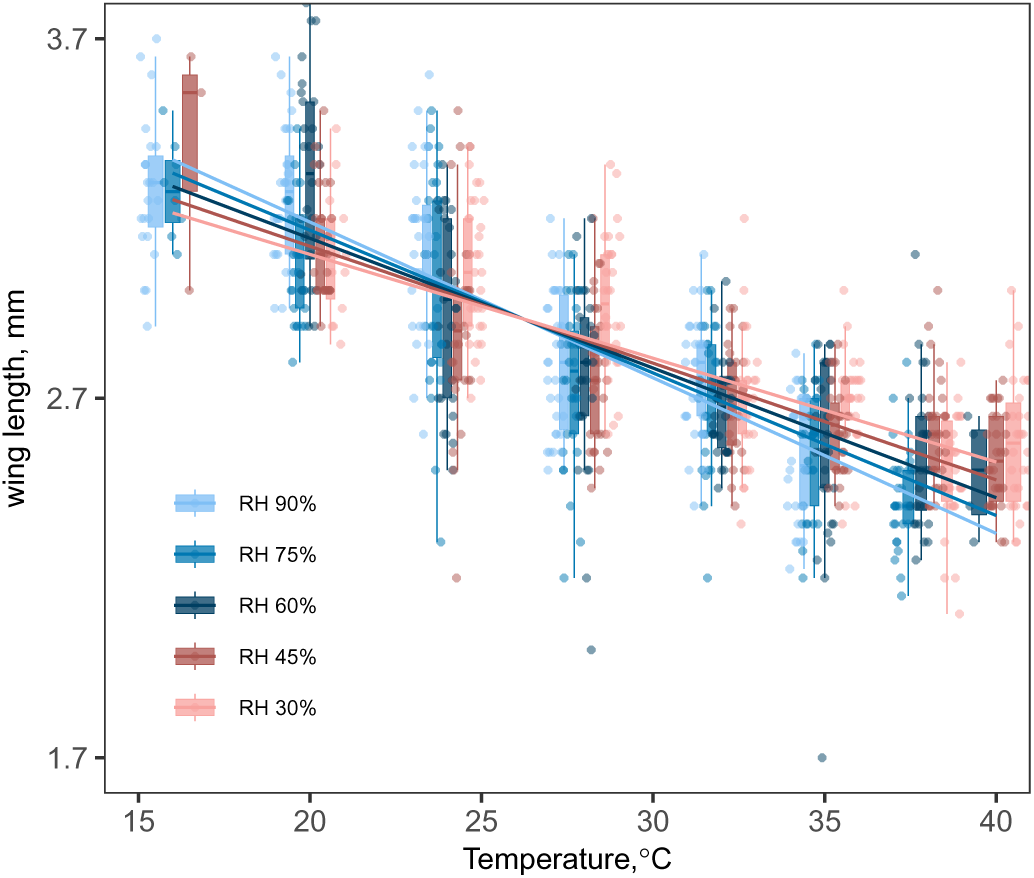
Relative air humidity and temperature interact to modulate the temperature–size relationship in *Anopheles stephensi*. Regression lines show that body size decreased with temperature, increased with humidity, and decreased more steeply with temperature at higher humidity. Boxplot horizontal lines represent medians; lower and upper hinges are the 25th and 75th percentiles. Upper whiskers extend from the hinge to the largest value no further than 1.5 × inter– quartile range (IQR) from the hinge. The lower whisker extends from the hinge to the smallest value at most 1.5 × IQR of the hinge. Points represent individual mosquitoes.

The interaction between temperature and humidity also caused significant variation in adult body size in the uncontrolled treatments (Fig. S5). Size decreased both at warmer temperatures (Slope: –0.10998; CI: –0.1397837, –0.0805072) and at lower humidity levels (Slope: –0.0301336; CI: –0.0412171, –0.0191704), but in contrast with the controlled treatments the decrease with temperature was greater in the lower humidity levels (75 and 60% RH) than at the highest humidity level (90%; Interaction: 0.0007560; CI: 0.0004171, 0.0010998). As temperatures increased from 16 to 40*^◦^*C, size was predicted to decrease by ∼1.43mm at 60% RH, whereas size at 90% RH decreased by ∼0.92mm.

### Relative humidity effects on temperature–dependent mosquito population growth rates

At all humidity levels, the maximal population growth rate (*r*_m_) exhibited a unimodal relationship with temperature, with *r*_m_ being positive above ∼15*^◦^*C and below ∼40*^◦^*C with optima between 0.27 and 0.29 (Fig. 3a–c; Table S5, S6). However, humidity systematically shifted the specific peak locations, levels, and upper and lower limits. For example, *T* _opt_ decreased by about 4.6*^◦^*C as RH levels increased from 30% to 90% ( Fig. 3b; Table S5). At the lowest humidity level (30%), *r*_m_ peaked at 36.09*^◦^*C, whereas, at the highest level (90%) it peaked at 31.48*^◦^*C. At intermediate humidity levels, *r*_m_ peaked between 33.48*^◦^*C and 34.48*^◦^*C (Fig. 3b; Table S5). The lowest *T*_min_ for *r*_m_ was predicted at 90% RH (12.46*^◦^*C), and it was predicted to be highest at 30% RH (16.43*^◦^*C;

**Figure 3:**
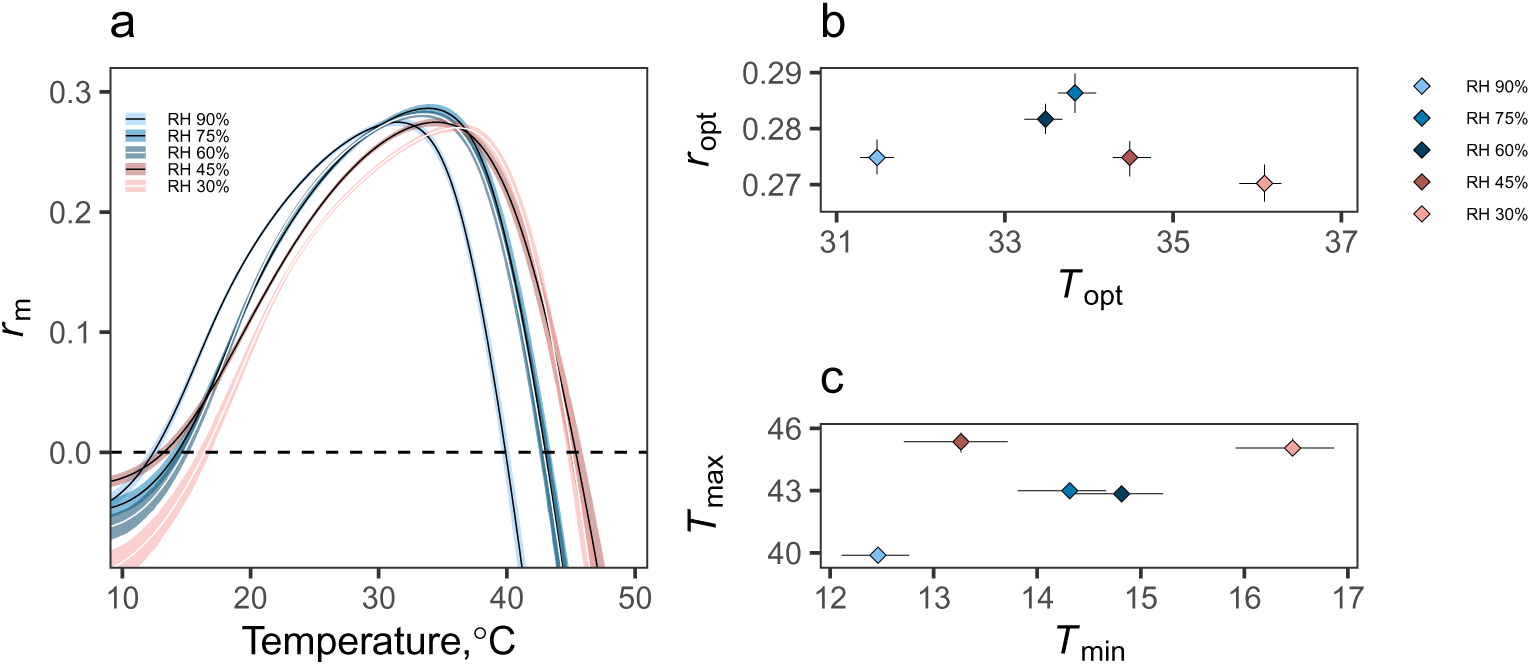
Effects of relative humidity on juvenile fitness traits shape the temperature dependence of maximal population growth rate. *r*_m_. (a–c) a. *r*_m_ TPCs across relative humidity levels. (b–c) b. *r*_opt_s versus *T*_opt_s, c. *T*_min_ versus *T*_max_ across humidity levels. Prediction bounds in **a** are HPD intervals calculated using the posteriors for each humidity–dependent TPC. Points (medians) in **b** and **c** were estimated numerically from the posterior distributions for each humidity level; bidirectional error bars are 95% credible intervals (CIs) calculated from the HPD intervals. Non– overlapping CIs indicate statistical significance to at least the *p* < 0.05 level in a classical analysis.

Fig. 3c; Table S6). At intermediate RH levels (45–75% RH), *T*_min_ was between 13.3*^◦^*C and 14.3*^◦^*C (Fig. 3c, Table S6). Similarly, *r*_m_ *T*_max_ was lowest at 90% RH (39.84*^◦^*C); it then increased to ∼43*^◦^*C at 75% and 60% RH. The highest *T*_max_ was predicted for the lowest humidity levels (45% and 30%) where it was ∼45*^◦^*C.

Humidity had a similar effect on *r*_opt_ in the uncontrolled treatments (Fig. S4, Tables S11 & S12). Predicted *r*_opt_ was between 0.27 and 0.30 at all humidity levels, and *T*_opt_ increased from ∼32*^◦^*C to ∼34*^◦^*C as RH decreased from 90% to 60%. Similarly, *r*_m_’s *T*_min_ was lowest when humidity was highest (∼11*^◦^*C at 90% RH) and it was highest at the lowest RH level (60%) where it was ∼26*^◦^*C. Predicted *r*_m_ *T*_max_ for RH levels above 60% in the uncontrolled treatments was between ∼40*^◦^*C and ∼42*^◦^*C, which was similar to the predictions for this parameter in the controlled evaporation treatments (i.e., between ∼40*^◦^*C and ∼43*^◦^*C at RH levels *>*60%).

### Sensitivity analysis

Variation in relative humidity influenced the sensitivity of *r*_m_, defined here as the rate at which *r*_m_ changes with temperature at a given temperature, to juvenile survival and development time (Fig. 4). High humidity dampened the sensitivity of *r*_m_ to both temperature and juvenile survival below 25*^◦^*C, while increasing sensitivity above 30*^◦^*C. Although *r*_m_ was relatively insensitive to development time, relative humidity increased this sensitivity at warm temperatures, increasing the magnitude of their negative relationship. This underlines how the temperature dependence of *r*_m_ mainly derives from how humidity influences juvenile survival, which determines the number of reproducing individuals. Similar patterns were observed for uncontrolled treatments; however, the sensitivity was significantly higher at both extremes of temperature and the dampening effect of relative humidity at low temperatures was much more apparent (Fig. S8).

**Figure 4:**
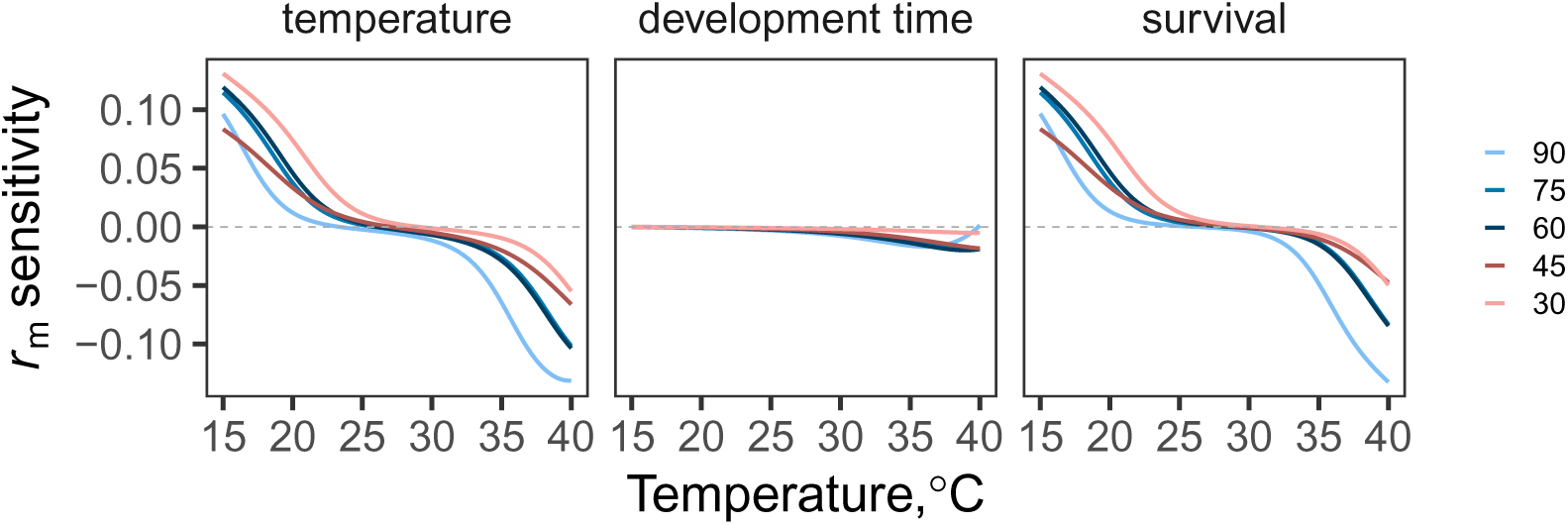
Sensitivity of maximal population growth rate, *r*_m_ to juvenile trait responses. Estimates of *r*_m_ are highly sensitive to temperature, driven almost entirely by sensitivity to juvenile survival, (*p*_EA_ in Eqn. 2, Table 2). This sensitivity is highest at the low and high temperature extremes, with relatively low sensitivity from 22*^◦^*C–30*^◦^*C. Relative humidity modulates the sensitivity of *r*_m_ to *p*_EA_, by increasing sensitivity at warmer temperatures and decreasing sensitivity at cooler temperatures. *r*_m_ shows relatively low sensitivity to juvenile development time (*α* in Eqn. 2, Table 2).

### Climatic suitability for mosquito population growth

Higher *r*_m_ values were predicted by the temperature–only humidity model in warmer, tropical regions, while lower values were predicted in in cooler higher–elevation areas including the Himalayas, Atlas Mountains, and Ethiopian Highlands (Fig. 5a, c). Incorporating humidity into the model reduced across some of Africa’s hottest and driest regions, such as parts of the Sahara and Sahel regions, southwest Africa, and the Horn of Africa. Conversely, incorporating humidity resulted in higher *r*_m_ across most of central and southern Africa, as well as portions of North Africa, with the most significant increases occurring in high–elevation regions. These spatial patterns remained consistent year–round. However, the areas where incorporating humidity increased *r*_m_ shifted northward from January through June and then southward again from July through December with the movement of the intertropical convergence zone and associated high rainfall and humidity (Fig. 5b). In South Asia, incorporating humidity generally reduced *r*_m_ across a substantial portion of central India but increased *r*_m_ in smaller portions of northern Pakistan, Bangladesh, southern India, and Sri Lanka throughout most of the year (Fig. 5d). During the humid monsoon season, however, including humidity led to increased *r*_m_ values across most of India, particularly central India. The temperature–only model indicated a larger area of suitable habitat than the temperature × humidity model. In South Asia, the temperature–only model identified approximately 0.4 million km^2^ more suitable area for mosquito growth than the temperature × humidity model (3.1 million km^2^ vs. 2.7 million km^2^). Similarly, in Africa, the temperature–only model identified an additional 0.8 million km^2^ of suitable land compared to the temperature × humidity model (25 million × vs. 24.2 million km^2^). In both Africa and South Asia, models incorporating humidity predicted higher climate suitability during rainy seasons, while temperature–only models indicated higher climate suitability during the drier seasons.

**Figure 5:**
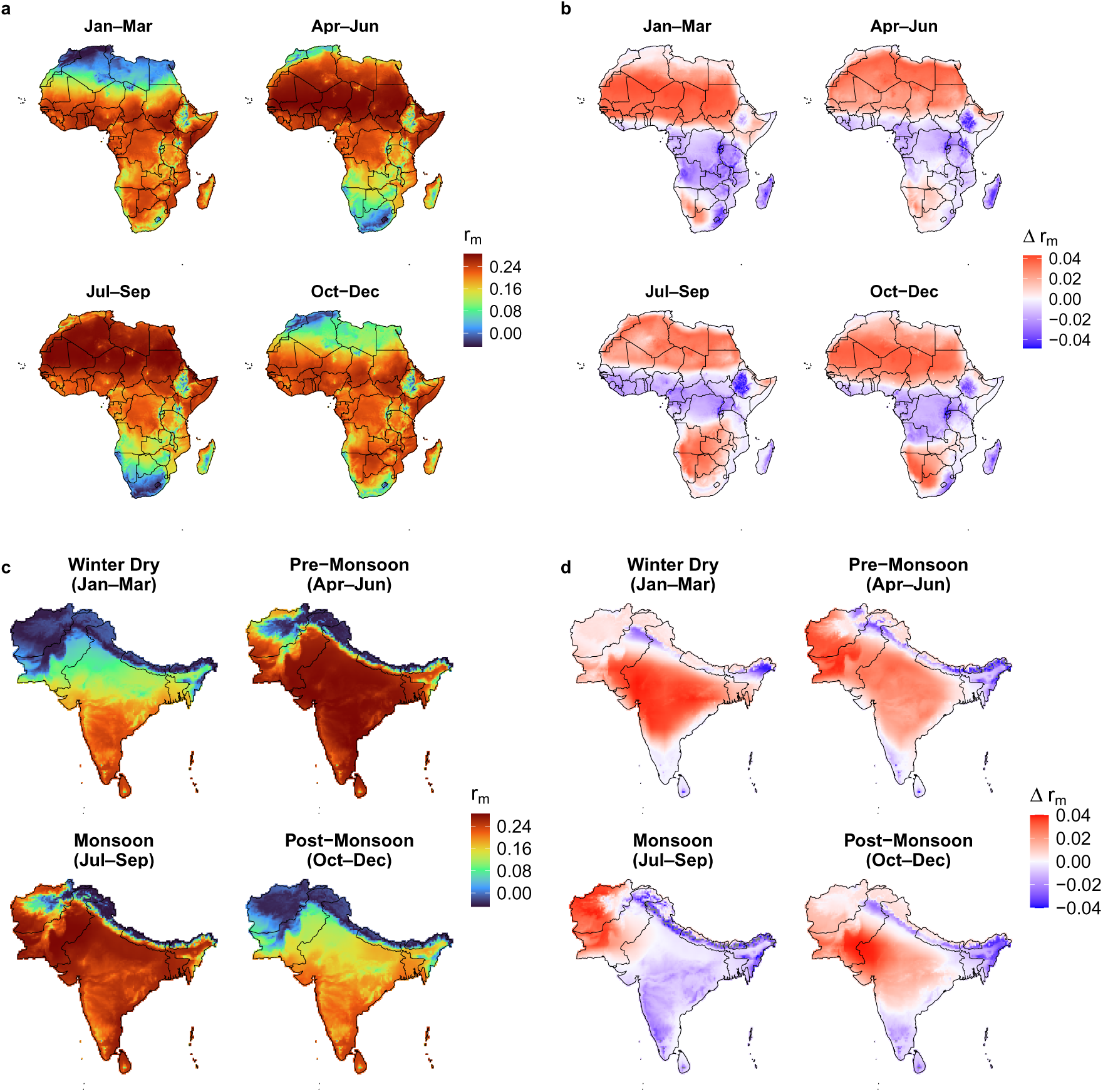
Seasonal spatial patterns of *r*_m_ derived from the NASA NEX-GDDP-CMIP6 daily historical dataset (1970–2000), for Africa (a and b) and South Asia (c and d) based on temperature and humidity. Maps (a) and (c) show the seasonal differences in *r*_m_ for the temperature-only model. In maps (b) and (d), Δ*r*_m_ = temperature-only *r*_m_ minus temperature- and humidity-dependent *r*_m_. Red (positive) values indicate that the temperature-only *r*_m_ model overestimated *r*_m_ and blue (negative) values indicate that temperature-only *r*_m_ model underestimated *r*_m_ (i.e., the temperature and humidity model predicted higher *r*_m_ than the temperature-only model). White values denote that both of the models were in agreement (Δ*r*_m_ = 0).

## Discussion and conclusion

Our results show that reductions in relative humidity during the aquatic developmental stages of *An. stephensi* systematically shift the temperature dependence of its maximal population growth rate, *r*_m_, toward warmer temperatures primarily via effects on juvenile survival and development. Peak juvenile survival (*T*_pk_) was ∼5*^◦^*C higher at the lowest humidity level (30%) than at the highest humidity level (90%), while the lower and upper thermal limits of this trait also differed to a similar extent (*T*_min_ was ∼16.5*^◦^*C at 30% RH and ∼12.5*^◦^*C at 90% RH; *T*_max_ was ∼40*^◦^*C at 30% RH and ∼45*^◦^*C at 90% RH) (Figs. 1a–c; Tables S1 & 2). Differences of this magnitude were not predicted for the development rate *T*_pk_s, which were all between ∼36*^◦^*C and ∼37.5*^◦^*C. However, the lower thermal threshold for this trait was ∼5*^◦^*C lower at 90% RH (8.63*^◦^*C) than at 30% RH (13.56*^◦^*C; Figs. 1d–e, Tables S3 & 4). Temperature × humidity interactions also significantly influenced adult body size. Size declined with increasing temperature across all humidity levels but it decreased more sharply with temperature at high humidity (e.g., ∼1mm at 90% RH vs. ∼0.07mm at 30% RH; Fig. 2). In treatments where evaporation could occur, these findings were more dramatic, with the steepest size declines in body size occurring at lower humidity levels (e.g., ∼1.43mm at 60% RH vs. ∼0.92mm at 90% RH; Fig. S5).

In our study, the effects of temperature we observed are consistent with previous research that has demonstrated non-linear and unimodal temperature-trait relationships in mosquito systems (Mordecai et al., 2019)—that is, temperatures towards the upper and lower limits constrained trait performance, whereas intermediate temperatures maximized it. Compared to similar studies on *An. stephensi*, the TPC parameter predictions from our intermediate RH levels for juvenile traits notably differ from estimates reported elsewhere in temperature–only studies. For example, survival peaked at ∼26*^◦^*C and development rate peaked at ∼32*^◦^*C in Villena et al. (2022), while, in the present study at intermediate humidity levels, these traits peaked at ∼29*^◦^*C and ∼36*^◦^*C, respectively. The predictions for other thermal performance parameters also differ between the two studies; in Villena et al. (2022), the *T*_min_s for juvenile survival and development rate were ∼18*^◦^*C and ∼19*^◦^*C, respectively. In contrast, in the present study, the *T*_min_s for these traits were ∼14*^◦^*C and ∼12*^◦^*C, respectively. This disagreement in the thermal performance of these traits likely stems from the way Villena et al. (2022) fitted TPCs to synthesized data on these traits from several studies that exposed their experimental populations to constant temperature ranges, but did not consider how variation in relative humidity across such ranges may have influenced their trait measurements. Despite differences in the extent to which body size decreased with temperature across humidity levels and evaporation settings, our results are consistent with the temperature-size rule in arthropods (Atkinson, 1995), including mosquitoes (Huxley et al., 2022; Agyekum et al., 2022), where adult body size decreased with developmental temperatures across all treatments.

Although the effects of temperature alone on juvenile mosquito traits are well-established, very little work has been done to understand the combined effects of variation in temperature and relative humidity on such traits (reviewed in Brown et al., 2023). We show that relative humidity variation did not alter the qualitative shape of temperature–trait relationships. However, it systematically shifted the overall thermal performance of juvenile survival and development rate toward warmer temperatures as relative humidity decreased. In addition, the strength of the negative relationship between temperature and adult body size upon emergence was also weaker under reduced humidity conditions and controlled evaporation. These results suggest that under humid conditions, juvenile development, survival, and body size upon emergence will be more constrained as temperatures warm compared to individuals developing in less humid environments. Such effects could have important implications for seasonal and spatial variation in the productivity of aquatic habitats, as well as mosquito fitness, mosquito pathogen competence, and population dynamics (Ameneshewa and Service, 1996; Moller-Jacobs et al., 2014; Shapiro et al., 2016). Evidence from the field on *Aedes albopictus* supports these results, with increases in daily mean relative humidity resulting in decreased adult emergence during the warm summer season (Murdock et al., 2017).

We propose several hypotheses that could describe how temperature and relative humidity interact to affect juvenile life history traits. However, we note that these hypotheses are not necessarily mutually exclusive. **First**, variation in temperature and humidity alters the surface tension of water (Pérez-Díaz et al., 2012). Higher temperatures increase the kinetic energy of water molecules, weakening hydrogen bonds and reducing surface tension. Increased humidity further lowers surface tension by reducing the net evaporation rate, as water vapor acts as a surfactant (Pérez-Díaz et al., 2012). Thus, we expect that in cool, dry environments, high surface tension could increase the metabolic demands of diving, foraging, and accessing atmospheric oxygen. Conversely, in hot, humid conditions, low surface tension may hinder successful emergence from the water (López-Doval et al., 2010). Supporting this idea, juvenile mosquito mortality has been shown to increase under surface tension extremes across a variety of species (Singh et al., 1957). **Second**, temperature and humidity influences oxygen concentrations in the air and water (Koue, 2024). Increases in temperature and humidity will decrease oxygen per unit volume due to air expansion and saturation with water molecules (Zhu et al., 2024; Shi et al., 2021), so, in cool, humid environments, atmospheric oxygen concentration may be higher, but high surface tension could impede access to it. In warm, humid environments, access to atmospheric oxygen will be easier, but its oxygen content will be less. As *Anopheles* juveniles breath atmospheric oxygen at the water surface (Mamai et al., 2016; Ha et al., 2017), limited access to it could constrain juvenile development, survival, metamorphosis, probability of successful emergence, and body size upon emergence (Briegel, 1990b; Clements, 1992; Paaijmans et al., 2008). Additionally, warmer water holds less dissolved oxygen, potentially further limiting access to oxygen affecting *An. stephensi* diving ability, feeding (Clements, 1992), and predator avoidance (Futami et al., 2008). However, work on other insect systems could indicate that for oxygen to substantially limit growth and survival directly, overall access to atmospheric oxygen would need to be substantially reduced to levels that may or may not be realistic for juvenile mosquitoes (Harrison and Haddad, 2011). **Third**, evaporative cooling may influence juvenile temperature exposure. In low humidity, water evaporates faster, cooling the surface more effectively (Brown et al., 2023) and increasing dissolved oxygen (Kurose et al., 2009; Fukatani et al., 2016). This could allow surface–residing larvae and pupae to survive and grow in spite of higher ambient temperatures. Resolving these mechanisms is critical to understanding how mosquito distributions and their population dynamics will respond to climate and land use change. These proposed mechanisms likely have broad implications for other aquatic organisms.

The effects of humidity on individual temperature–trait relationships varied, but it consistently influenced the maximal population growth rate, *r*_m_. Decreases in relative humidity shifted *r*_m_ *T*_opt_ and thermal limits (*T*_min_ and *T*_max_) toward warmer temperatures (Fig. 3, Tables S5, S6). However, the peak value of *r*_m_ at *T*_pk_ (*r*_opt_) remained relatively stable, indicating that humidity primarily affects the horizontal position of the *r*_m_ curve, not its shape. While temperature interactions with biotic factors like resource competition are known to influence arthropod populations (Park, 1954; Mueller, 1988), including mosquitoes (Huxley et al., 2022; Evans et al., 2021), mechanistic studies on abiotic interactions—particularly temperature and humidity—are rare (Brown et al., 2023). Sensitivity analyses (Figs. 4, S8) show that humidity–driven changes in the temperature dependence of juvenile survival (*p*_EA_ in Eqn. 2) largely explain shifts in the temperature dependence of *r*_m_. At both low and high temperatures, humidity modulated the impact of juvenile survival on *r*_m_, while juvenile development time (*α* in Eqn. 2) showed relatively low sensitivity. Specifically, population–level reproductive output decreased when both temperature and humidity were high due to fewer individuals reaching maturity, underscoring survival as the primary driver of humidity’s effect on temperature-dependent population growth (Figs. 1, 3). The relative importance of juvenile survival in shaping the temperature dependence of *r*_m_ is further underlined by the lower and upper thermal limits (*T*_min_ and *T*_max_, respectively; Fig. 1, Table S2) predicted for this trait.

As noted above, this trait’s *T*_min_ was about 4*^◦^*C higher and its *T*_max_ was ∼5*^◦^*C higher at the lowest humidity level (30% RH) compared to the highest humidity level (90% RH). It is this shift in the temperature dependence of juvenile survival that was mainly responsible for the similar shift predicted for *r*_m_’s *T*_min_ and *T*_max_ (Fig. 3a & c).

The inclusion of relative humidity in thermal performance models for *An. stephensi* led to distinct historical patterns of climate suitability for population growth in India and Africa compared to the temperature–only model (Fig. 5; see Fig. S9 for maps showing seasonal mean temperatures and relative humidity for 1970–2000 across Africa and India). For example, the temperature– only model predicted larger areas of climatic suitability in South Asia (0.4 million km^2^ more) and Africa (0.8 million km^2^ more) to be climatically suitable for *An. stephensi* compared to models that included both temperature and relative humidity. The largest discrepancies occurred in hotter regions. Incorporating relative humidity increased the thermal suitability in warm, tropical zones but reduced suitability in hot, dry areas, with the exception of higher elevations. Seasonal differences in climate suitability were also evident when accounting for variation in temperature and humidity across winter (cool, dry), pre–monsoon (hot, dry), monsoon (warm, humid), and post-monsoon (cool, moderate humidity) periods in India. Models incorporating humidity predicted higher suitability during rainy seasons, whereas temperature–only models predicted greater suitability in the other seasons that were drier. Similar trends are observed for Africa. These results reflect that mean temperatures across both India and Africa typically remain below the predicted thermal optima for *r*_m_ regardless of relative humidity. Importantly, if climate warming, intensifying urban heat island effects, and heat extremes during the hot seasons push temperatures above ∼35*^◦^*, *An. stephensi* suitability may remain high if relative humidity concurrently drops. Overall, our modeling and maps highlight the importance of including both temperature and humidity when projecting the impacts of environmental change on vector–borne disease dynamics and other arthropod–mediated systems.

The results of our study should be interpreted in light of several limitations. First, the experiment was conducted under laboratory conditions and does not account for field–relevant variation such as diurnal environmental fluctuations (Shocket et al., 2025), locally adapted mosquito populations (Dennington et al., 2024; Couper et al., 2025; Sternberg and Thomas, 2014), and behavioral responses to sub–optimal climates (Verhulst et al., 2020). Nonetheless, this study provides a foundational step toward understanding how relative humidity can affect the temperature dependence of mosquito fitness and their population dynamics. Our findings align with field studies in other vectors (e.g., *Ae. albopictus*; Murdock et al., 2017; Evans et al., 2019) and epidemiological work in the *An. stephensi*-malaria system (Santos-Vega et al., 2022). Second, our metric of climate suitability, *r*_m_, excludes effects of temperature or relative humidity on adult traits such as mortality and fecundity. Temperature effects on these traits are likely influenced by humidity directly (Brown et al., 2023), as well as potentially through carry-over effects on body size and condition at emergence (Moller-Jacobs et al., 2014; Shapiro et al., 2016). Third, environmental suitability also depends on the availability of aquatic habitat for juvenile development, which is shaped by precipitation, piped water access, storage practices, and evaporation (Stewart Ibarra et al., 2013; Hayden et al., 2010; Whittaker et al., 2023). While *An. stephensi* often develops in larger water containers that are less prone to evaporation (Thomas et al., 2016), other vectors like *Aedes spp.* use smaller, ephemeral habitats more sensitive to drying (Day, 2016). Further, while the effects of relative humidity on juvenile development, survival, and population growth rate were similar, we did see differences in the effect on adult body size suggesting mechanisms shaping potential carry–over effects could vary in environments subject to evaporation. Finally, although we controlled the initial amount of resources juvenile mosquitoes had access to, variation in temperature and humidity conditions could affect microbial growth, which combined with effects on juvenile survival, could alter the effects of resource competition on *r*_m_ and the efficacy of vector control strategies (Guégan et al., 2018; Coon et al., 2016; Takken et al., 2013; White et al., 2011).

In conclusion, our findings challenge current paradigms in mosquito thermal biology by highlighting the significant role of relative humidity. We show that humidity strongly influences the thermal performance of aquatic life stages in *An. stephensi*, a key malaria vector. Lower humidity levels shifted optimal temperatures for development, survival, and population growth toward warmer temperatures, with broad implications for predicting mosquito distributions. These results suggest that temperature and humidity together may drive local adaptation and shape mosquito responses to environmental change. Importantly, variation in these factors could alter the effectiveness, reach, and cost of larval control programs. Future research should examine how humidity affects adult traits and malaria transmission, validate findings under field conditions, and model population dynamics under projected climate and land-use changes. *An. stephensi* exhibited high phenotypic plasticity in response to variation in temperature and relative humidity, which will likely facilitate its ability to establish and persist in new regions, as well as to adjust to future environmental change. Ultimately, this work advances our understanding of how multiple climate variables shape mosquito ecology, with broader relevance to the conservation and risk assessment of other temperature–sensitive species.

## Data and code availability

All data and code needed to evaluate the conclusions in the paper are present in the paper, the Supplementary Materials, or the project’s GitHub repository found at:

