## Supplementary Material for "Beyond temperature: Relative humidity systematically shifts the temperature dependence of population growth in a malaria vector"

#### Supplementary Equations

##### *Modelling the development time TPCs*

As noted in the main text, we modelled the development time TPCs for each humidity level in two ways. First, to get the juvenile development time ($\alpha$) for our $r_m$ calculations (Eqn. 2, Main text), we implemented an exponential decay function (SE1) in the `bayesTPC` package in R; Sorek et al., 2025) to directly fit to the observed data for this trait (Figure S1)

$$\alpha(T) = a^{(-rT)} + c \tag{SE1}$$

Here, $a$ is a scaling factor determining the amplitude of the decay, $r$ is the rate of decay (where $r$ is $> 0$), $T$ is the temperature in degrees Celsius, and $c$ is a constant that represents the baseline that $\alpha(T)$ approaches as $T \to \infty$.

However, this model (SE1) is not unimodal and does not have defined values for $T_{\text{min}}$, $T_{\text{pk}}$, $B_{\text{pk}}$, and $T_{\text{max}}$. Thus in order to be able to make a more direct comparison to previous studies, we fitted the standard Briere model (SE2; as implemented in the `bayesTPC` package in R; Briere et al., 1999; Sorek et al., 2025) to *inverted development times* (i.e., development rate; $1/\alpha$ in Eqn. 2).

$$1/\alpha(T) = q \cdot T \cdot (T - T_{\text{min}}) \cdot \sqrt{(T_{\text{max}} > T) \cdot (T_{\text{max}} - T)} \cdot (T_{\text{max}} > T) \cdot (T > T_{\text{min}}) \tag{SE2}$$

Here, $T$ is temperature in degrees Celsius, $T_{\text{min}}$ is the low temperature (ºC) at which rates become negative, $T_{\text{max}}$ is the high temperature (ºC) at which rates become negative, and $q$ scale parameter that sets the maximum rate of the curve. The temperatures at which trait performance peaks and its value at its peak ($T_{\text{pk}}$ and $B_{\text{pk}}$, respectively; Table 2, Main text) were estimated numerically from the posterior distributions for each humidity level's TPC and summarized by the posterior medians and Highest Posterior Density (HPD) intervals.

##### *Modelling the survival probability TPCs*

We parameterized $p_{\text{EA}}$ in Eqn. 2 (Main text) by substituting TPCs fitted to the binomial juvenile survival data for each humidity level. We fitted to these data using the binomial quadratic glm model that is implemented in the R package, `bayesTPC` (Sorek et al., 2025):

$$p_{\text{EA}}(T) = b0 + b1 \cdot T + (b2 \cdot T^2) \tag{SE3}$$

Here, $T$ is temperature in degrees Celsius, $b_0$ is the value of $p_{\mathrm{EA}}$ when $T$ is zero and $b_1$ determines the linear realtionship between temperature and survival probability. In the quadratic term, $b_2 \cdot T^2$, $b_2$ models the relationship's curvature.

#### Trait sensitivity analysis

To determine the extent to which variation in humidity can affect the relative contributions of the individual juvenile fitness traits to $r_{\mathrm{m}}$'s temperature dependence, using the chain rule, we can write (Cator et al., 2020; Mordecai et al., 2013):

$$\frac{\mathrm{d}r_{\mathrm{m}}}{\mathrm{d}T} = \frac{\partial r_{\mathrm{m}}}{\partial b_{\mathrm{max}}}\frac{\mathrm{d}b_{\mathrm{max}}}{\mathrm{d}T} + \frac{\partial r_{\mathrm{m}}}{\partial \alpha}\frac{\mathrm{d}\alpha}{\mathrm{d}T} + \frac{\partial r_{\mathrm{m}}}{\partial z}\frac{\mathrm{d}z}{\mathrm{d}T} + \frac{\partial r_{\mathrm{m}}}{\partial p_{\mathrm{EA}}}\frac{p_{\mathrm{EA}}}{\mathrm{d}T} + \frac{\partial r_{\mathrm{m}}}{\partial \kappa}\frac{\mathrm{d}\kappa}{\mathrm{d}T}. \tag{SE4}$$

Each summed term in the right hand side of this equation quantifies the relative contribution of each trait TPC parameter in Eqn. 2 (Main text) to the temperature dependence of $r_{\mathrm{m}}$. To calculate the derivatives for each TPC parameter in SE4, we used the MAP (Maximum A Posteriori) estimator for each trait–humidity combination (Figs. 4 & S8). The sample-based MAP estimator is calculated as part of the MCMC fitting process in the `bayesTPC` package (Sorek et al., 2025) in R. We used this method to estimate the Eqn. 2 parameters because it maximizes the unknown parameter's posterior probability, given the observed data and prior beliefs so, compared to the Maximum Likelihood Estimator (MLE), it incorporates prior information.

#### Methods for Mapping Study Region and Data Sources

We mapped the maximal mosquito population growth rate ($r_{\mathrm{m}}$) across South Asia and Africa under historical and future climate conditions. The South Asian study region encompassed India and its neighboring countries (Bangladesh, Bhutan, Nepal, Sri Lanka, Pakistan, Maldives, and Afghanistan), capturing diverse climates from tropical to temperate zones. The African study region covered the entire continent, incorporating equatorial, savannah, arid, and subtropical climates to enable broad regional comparisons of potential mosquito growth.

Climate variables, including daily maximum temperature, minimum temperature, and specific humidity, were obtained from the NASA NEX–GDDP–CMIP6 dataset. These statistically downscaled, bias–corrected projections have a resolution of $0.25^\circ \times 0.25^\circ$ ($\sim$27.8 km). We incorporated outputs from 23 general circulation models (GCMs) in this dataset, analyzing the ensemble of historical (1970–2000) model runs to generate climate inputs for the models.

##### *Climate Data Processing*

We used the NASA climate data to estimate ($r_{\mathrm{m}}$), which integrates the effects of temperature and humidity on mosquito population dynamics. Daily climate variables including maximum temperature, minimum temperature, and specific humidity from 1970 to 2000 were masked to terrestrial grid cells within the study–region boundaries. These daily data were first aggregated into monthly averages. Monthly mean temperatures were calculated as the average of daily maximum and minimum temperatures. Relative humidity was computed from monthly averaged specific humidity data using the Magnus–Tetens equation, accounting for temperature-dependent saturation vapor pressure. Finally, seasonal means were calculated by averaging monthly data across defined three–month periods (e.g., January—March) for the entire 1970–2000 period.

##### *Trait-Based Population Growth Model*

Monthly mosquito population growth was predicted using the trait-based model of the intrinsic rate of population increase ($r_{\mathrm{m}}$). Biological traits, including juvenile survival, developmental rate, and fecundity, were modeled as nonlinear functions of temperature based on laboratory experiments. Temperature-trait curves were derived the experimental RH values of 30%, 45%, 60%, 75%, and 90%. To quantify the influence of humidity, we conducted two model runs. The temperature-only model held humidity constant at a fixed reference value of 75% RH, reflecting the baseline condition under which most temperature-trait models are derived. The temperature $\times$ RH model used the actual RH values from the climate dataset. We applied piecewise linear interpolation to handle continuous RH variations in the climate data, deriving each modeled trait from the two nearest experimental RH levels. This method ensured smooth trait responses to varying humidity without abrupt category transitions. These analyses produced monthly raster datasets of $r_{\mathrm{m}}$ values for the two model runs, and the monthly outputs were summarized to generate mean annual $r_{\mathrm{m}}$.

##### *Spatial Analysis and Mapping*

Maps of mean annual $r_{\mathrm{m}}$ were generated to highlight spatial variability in mosquito population growth potential. Difference maps ($\Delta r_{\mathrm{m}}$) were created to highlight areas where adding humidity altered the growth rates, revealing regions of higher or lower suitability for population growth. We also computed the total land area for which climate was suitable for mosquito growth ($r_{\mathrm{m}} > 0$) during all months of the year (Fig. S9).

#### Supplementary figures and tables

**Table S1: Estimates for the juvenile survival TPC parameters; $B_{pk}$ and $T_{pk}$.** Numerical posterior estimates (median ± 95% credible intervals) of the parameters from $n$ = 10,000 iterations for a burn–in of $n$ = 5000.

| RH (%) | Bpk median | Bpk lower | Bpk upper | Tpk median | Tpk lower | Tpk upper |
|---|---|---|---|---|---|---|
| 90 | 0.937 | 0.924 | 0.949 | 26.176 | 26.026 | 26.326 |
| 75 | 0.918 | 0.903 | 0.931 | 28.629 | 28.478 | 28.829 |
| 60 | 0.901 | 0.885 | 0.917 | 28.829 | 28.629 | 29.029 |
| 45 | 0.835 | 0.816 | 0.857 | 29.279 | 29.079 | 29.530 |
| 30 | 0.893 | 0.876 | 0.910 | 30.731 | 30.531 | 30.931 |

**Table S2: Estimates for the juvenile survival TPC parameters; $T_{min}$ and $T_{max}$.** Numerical posterior estimates (median ± 95% credible intervals) of the parameters from $n$ = 10,000 iterations for a burn–in of $n$ = 5000.

| RH (%) | Tmin | Tmin lower | Tmin upper | Tmax | Tmax lower | Tmax upper |
|---|---|---|---|---|---|---|
| 90 | 12.312 | 11.962 | 12.663 | 39.990 | 39.640 | 40.240 |
| 75 | 14.164 | 13.714 | 14.565 | 43.093 | 42.693 | 43.493 |
| 60 | 14.715 | 14.164 | 15.065 | 42.993 | 42.593 | 43.343 |
| 45 | 13.063 | 12.563 | 13.564 | 45.495 | 45.045 | 45.996 |
| 30 | 16.316 | 15.766 | 16.717 | 45.195 | 44.745 | 45.596 |

**Table S3: Estimates for the juvenile development rate TPC parameters; $B_{pk}$ and $T_{pk}$.** Numerical posterior estimates (median ± 95% credible intervals) of the parameters from $n$ = 10,000 iterations for a burn–in of $n$ = 5000.

| RH (%) | Bpk median | Bpk lower | Bpk upper | Tpk median | Tpk lower | Tpk upper |
|---|---|---|---|---|---|---|
| 90 | 0.141 | 0.135 | 0.147 | 36.236 | 34.885 | 37.037 |
| 75 | 0.144 | 0.140 | 0.148 | 36.436 | 35.536 | 37.137 |
| 60 | 0.150 | 0.144 | 0.156 | 37.187 | 36.737 | 37.538 |
| 45 | 0.142 | 0.139 | 0.145 | 37.337 | 36.887 | 37.487 |
| 30 | 0.132 | 0.125 | 0.138 | 37.437 | 36.887 | 37.788 |

**Table S4: Estimates for the juvenile development rate TPC parameters; $T_{min}$ and $T_{max}$.** Posterior estimates (mean ± 95% credible intervals) of the standard Briere model (SE2) parameters from $n$ = 10,000 iterations for a burn–in of $n$ = 5000.

| RH (%) | Tmin | Tmin lower | Tmin upper | Tmax | Tmax lower | Tmax upper |
|---|---|---|---|---|---|---|
| 90 | 8.630 | 6.690 | 10.647 | 43.916 | 42.290 | 45.00 |
| 75 | 11.289 | 9.734 | 12.695 | 43.845 | 42.802 | 44.99 |
| 60 | 11.418 | 9.210 | 13.722 | 44.736 | 44.249 | 45.00 |
| 45 | 12.420 | 11.348 | 13.409 | 44.745 | 44.321 | 45.00 |
| 30 | 13.562 | 11.787 | 14.996 | 44.685 | 44.087 | 45.00 |

**Table S5: Estimates for the $r_m$ TPC parameters; $T_{opt}$ and $r_{opt}$.** Numerical posterior estimates (median ± 95% credible intervals) of the $r_m$ model (Eqn. 1, Main text) parameters from $n$ = 10,000 iterations for a burn–in of $n$ = 5000.

| RH (%) | Topt | Topt lower | Topt upper | ropt median | ropt lower | ropt upper |
|---|---|---|---|---|---|---|
| 90 | 31.481 | 31.281 | 31.682 | 0.275 | 0.272 | 0.278 |
| 75 | 33.834 | 33.634 | 34.084 | 0.286 | 0.283 | 0.290 |
| 60 | 33.483 | 33.233 | 33.684 | 0.282 | 0.279 | 0.284 |
| 45 | 34.484 | 34.284 | 34.735 | 0.275 | 0.271 | 0.278 |
| 30 | 36.086 | 35.786 | 36.286 | 0.270 | 0.267 | 0.274 |

**Table S6: Estimates for the $r_m$ TPC parameters; $T_{min}$ and $T_{max}$.** Numerical posterior estimates (median ± 95% credible intervals) of the parameters from $n$ = 10,000 iterations for a burn–in of $n$ = 5000.

| RH (%) | Tmin | Tmin lower | Tmin upper | Tmax | Tmax lower | Tmax upper |
|---|---|---|---|---|---|---|
| 90 | 12.462 | 12.112 | 12.763 | 39.890 | 39.590 | 40.140 |
| 75 | 14.314 | 13.814 | 14.665 | 42.993 | 42.593 | 43.393 |
| 60 | 14.815 | 14.364 | 15.215 | 42.843 | 42.492 | 43.193 |
| 45 | 13.263 | 12.713 | 13.714 | 45.345 | 44.845 | 45.796 |
| 30 | 16.466 | 15.916 | 16.867 | 45.045 | 44.645 | 45.495 |

**Figure S1: Observed juvenile survival data (hatching–to–adult) across temperature–humidity levels.** There were $n$=3 replicates per treatment each containing $n$=100 L1 larvae at the start of the experiment. The figure shows the number of survivors for each treatment from a pooled total of $n$=300 at the start of the experiment. No individuals survived to adulthood at 14°C and 42°C irrespective of humidity level.

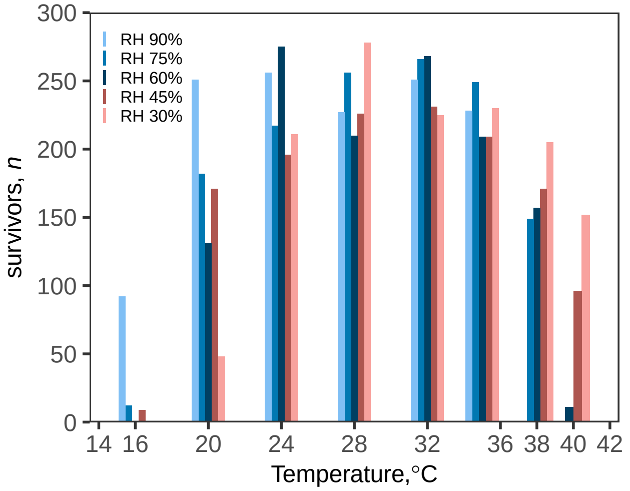


**Figure S2: Juvenile development time TPCs (hatching–to–adult) used for the temperature– and humidity–dependent $r_m$ calculations.** Development time ($\alpha$ in Eqn. 1, Main text) TPCs were fitted using Eqn. SE3. Points are individual mosquitoes. Relative humidity (%) levels are shown in the title boxes.

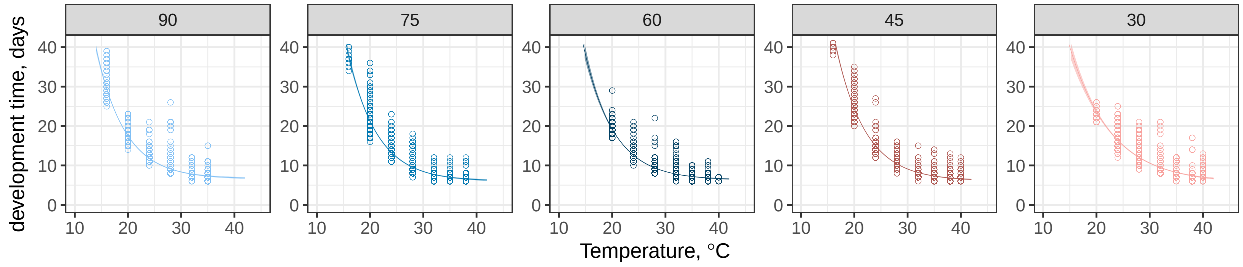

### *Results from uncontrolled evaporation experiments*

**Figure S3: Uncontrolled evaporation: Relative humidity shapes the temperature dependence of juvenile fitness traits in** *An. stephensi* **(a–f)**. a. Humidity–dependent survival probability TPCs ($p_{EA}$ in Eqn. 2, Main text). b–c. **b.** Numerical survival probability parameter estimates of $T_{min}$ and $T_{max}$ (Tables S7, S8). **c.** Predicted peak survival probabilities at $T_{pk}$ at each humidity level. Legend in b also applies to c. **d.** Humidity–dependent development rate TPCs ($1/\alpha$ in Eqn. 2). **e.** Development rate parameter estimates for $T_{min}$ versus $T_{max}$ across humidity levels (Table S10). **f.** Predicted peak development rate at $T_{pk}$ at each humidity level (Table S9). Prediction bounds in **a** and **d** are HPD intervals estimated from the posteriors for each TPC. In b–c and e–f, errors bars are 95% credible intervals (CIs). Non-overlapping CIs indicate statistical significance to at least the $p < 0.05$ level in a classical analysis.

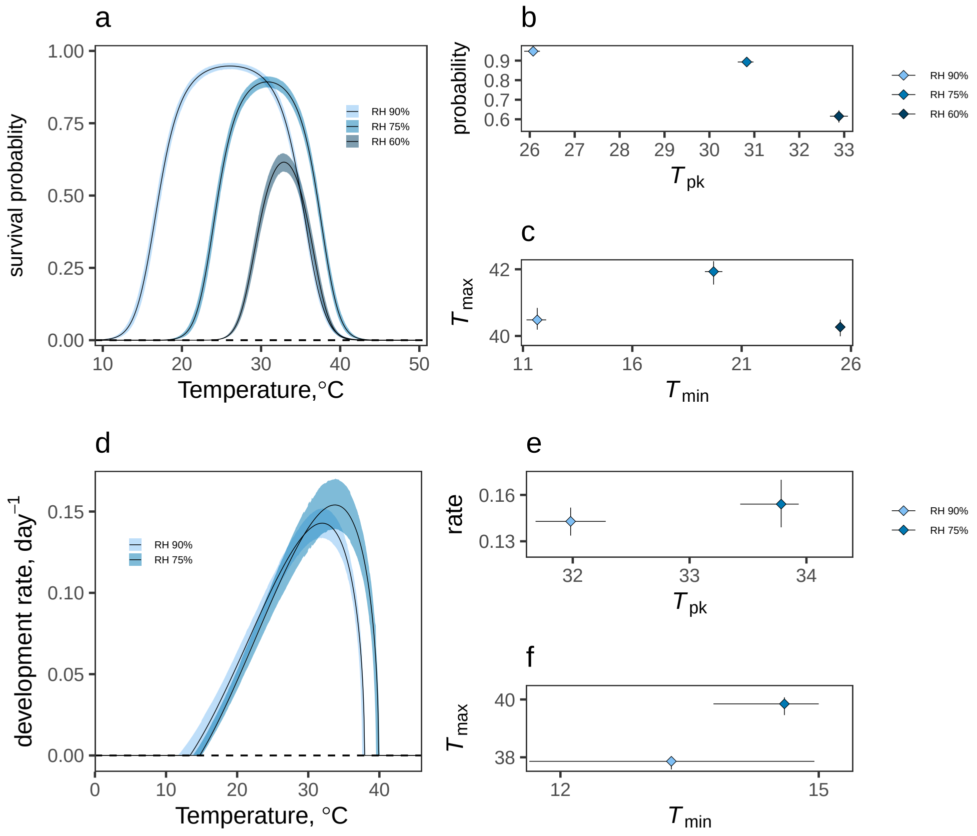

**Figure S4: Uncontrolled evaporation: Effects of relative air humidity on juvenile fitness traits shape the temperature dependence of maximal population growth rate, $r_m$.** (a–c) **a.** $r_m$ TPCs across relative humidity levels. (b–c) **b.** $r_{opt}$s versus $T_{opt}$s (Table S11), **c.** $T_{min}$ versus $T_{max}$ across humidity levels (Table S12). Prediction bounds in **a** are HPD intervals calculated using the posteriors for each humidity–dependent TPC. Error bars in **b** and **c** are 95% credible intervals (CIs). Non-overlapping CIs indicate statistical significance to at least the $p < 0.05$ level in a classical analysis.

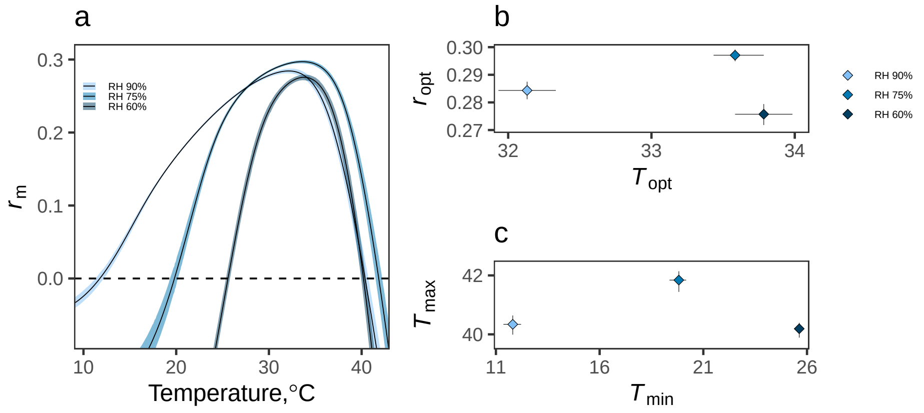


**Figure S5: Uncontrolled evaporation: Relative air humidity and temperature interact to modulate the temperature–size relationship in** ***Anopheles stephensi***. Regression lines show that body size decreased with temperature (Slope: –0.10998; CI: –0.1397837, –0.0805072) and humidity (Slope: –0.0301336; CI: –0.0412171, –0.0191704), and it decreased more steeply with temperature at lower humidity (Interaction: 0.0007560; CI: 0.0004171, 0.0010998). Boxplot horizontal lines represent medians; lower and upper hinges are the 25th and 75th percentiles. Upper whiskers extend from the hinge to the largest value no further than 1.5 × inter-quartile range (IQR) from the hinge. The lower whisker extends from the hinge to the smallest value at most 1.5 × IQR of the hinge. Points represent individual mosquitoes.

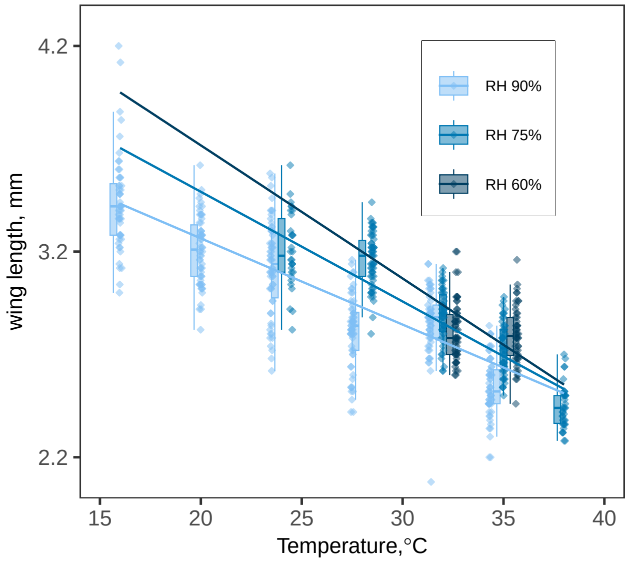

**Table S7: Estimates for the juvenile survival TPC parameters; $B_{pk}$ and $T_{pk}$.** Numerical posterior estimates (median ± 95% credible intervals) of the parameters from $n$ = 10,000 iterations for a burn–in of $n$ = 5000.

| RH (%) | Bpk median | Bpk lower | Bpk upper | Tpk median | Tpk lower | Tpk upper |
|---|---|---|---|---|---|---|
| 90 | 0.948 | 0.937 | 0.958 | 26.076 | 25.876 | 26.226 |
| 75 | 0.893 | 0.874 | 0.912 | 30.831 | 30.631 | 30.981 |
| 60 | 0.616 | 0.583 | 0.646 | 32.883 | 32.683 | 33.083 |

**Table S8: Estimates for the juvenile survival TPC parameters; $T_{min}$ and $T_{max}$.** Numerical posterior estimates (median ± 95% credible intervals) of the parameters from $n$ = 10,000 iterations for a burn–in of $n$ = 5000.

| RH (%) | Tmin | Tmin lower | Tmin upper | Tmax | Tmax lower | Tmax upper |
|---|---|---|---|---|---|---|
| 90 | 11.662 | 11.161 | 12.062 | 40.490 | 40.190 | 40.841 |
| 75 | 19.720 | 19.319 | 20.120 | 41.942 | 41.542 | 42.242 |
| 60 | 25.526 | 25.275 | 25.676 | 40.290 | 39.990 | 40.490 |

**Table S9: Estimates for the juvenile development rate TPC parameters; $B_{pk}$ and $T_{pk}$.** Numerical posterior estimates (median ± 95% credible intervals) of the parameters from $n$ = 10,000 iterations for a burn–in of $n$ = 5000.

| RH (%) | Bpk median | Bpk lower | Bpk upper | Tpk median | Tpk lower | Tpk upper |
|---|---|---|---|---|---|---|
| 90 | 0.143 | 0.134 | 0.152 | 31.982 | 31.682 | 32.282 |
| 75 | 0.154 | 0.139 | 0.170 | 33.784 | 33.433 | 33.934 |

**Table S10: Estimates for the juvenile development rate TPC parameters; $T_{min}$ and $T_{max}$.** Posterior estimates (mean ± 95% credible intervals) of the standard Briere model (SE2) parameters from $n$ = 10,000 iterations for a burn–in of $n$ = 5000.

| RH (%) | Tmin | Tmin lower | Tmin upper | Tmax | Tmax lower | Tmax upper |
|---|---|---|---|---|---|---|
| 90 | 13.288 | 11.638 | 14.95 | 37.861 | 37.579 | 38.008 |
| 75 | 14.601 | 13.777 | 15.00 | 39.849 | 39.460 | 40.074 |

**Table S11: Estimates for the $r_m$ TPC parameters; $T_{opt}$ and $r_{opt}$.** Numerical posterior estimates (median ± 95% credible intervals) of the parameters from $n$ = 10,000 iterations for a burn–in of $n$ = 5000.

| RH (%) | Topt | Topt lower | Topt upper | ropt median | ropt lower | ropt upper |
|---|---|---|---|---|---|---|
| 90 | 32.132 | 31.932 | 32.332 | 0.284 | 0.281 | 0.288 |
| 75 | 33.584 | 33.433 | 33.784 | 0.297 | 0.295 | 0.299 |
| 60 | 33.834 | 33.584 | 33.984 | 0.276 | 0.272 | 0.279 |

**Table S12: Estimates for the $r_m$ TPC parameters; $T_{min}$ and $T_{max}$.** Numerical posterior estimates (median ± 95% credible intervals) of the parameters from $n$ = 10,000 iterations for a burn–in of $n$ = 5000.

| RH (%) | Tmin | Tmin lower | Tmin upper | Tmax | Tmax lower | Tmax upper |
|---|---|---|---|---|---|---|
| 90 | 11.812 | 11.361 | 12.212 | 40.340 | 40.040 | 40.641 |
| 75 | 19.820 | 19.419 | 20.220 | 41.842 | 41.491 | 42.142 |
| 60 | 25.626 | 25.375 | 25.776 | 40.190 | 39.890 | 40.390 |

**Figure S6: Uncontrolled evaporation: Observed juvenile survival data (hatching–to–adult) across temperature–humidity levels.** There were $n$=3 replicates per treatment each containing $n$=100 L1 larvae at the start of the experiment. The figure shows the number of survivors for each treatment from a pooled total of $n$=300 at the start of the experiment. No individuals survived to adulthood at 14°C and 42°C irrespective of humidity level.

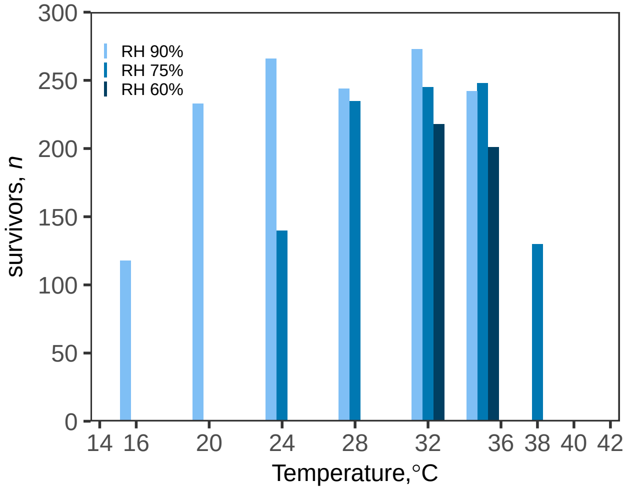

**Figure S7: Uncontrolled evaporation: Juvenile development time TPCs (hatching–to–adult) used for the temperature– and humidity–dependent $r_{\mathrm{m}}$ TPCs.** Development time ($\alpha$ in Eqn. 2) TPCs were fitted using Equation S1. Points are individual mosquitoes.

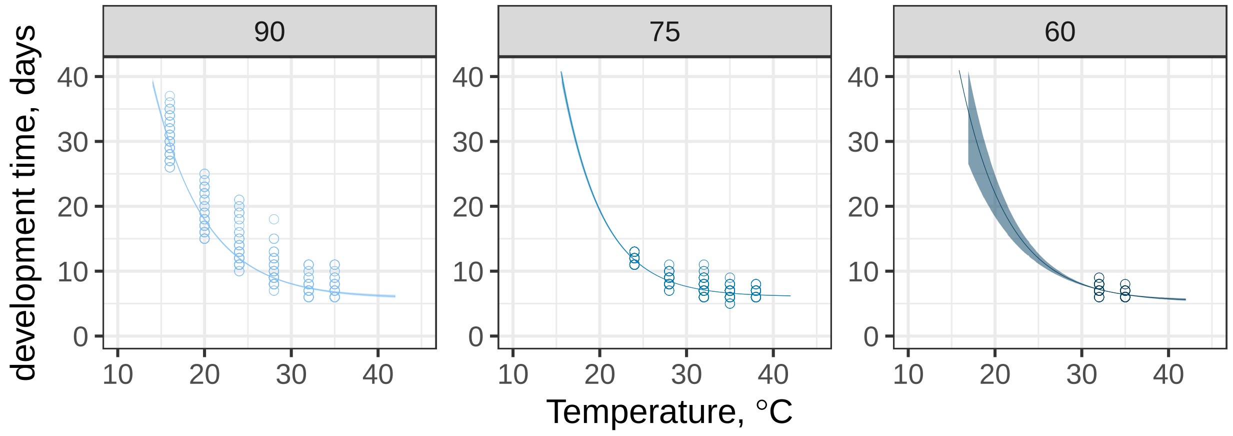


**Figure S8: Uncontrolled evaporation: Sensitivity of $r_{\mathrm{m}}$ to key parameters**. Similarly to the evaporation control experiment, $r_{\mathrm{m}}$ is highly sensitive to temperature, due to its effect on juvenile survival, $p_{\mathrm{EA}}$. However, sensitivity to temperature is much higher at low temperatures, and the buffering effect of relative humidity is much more noticeable.

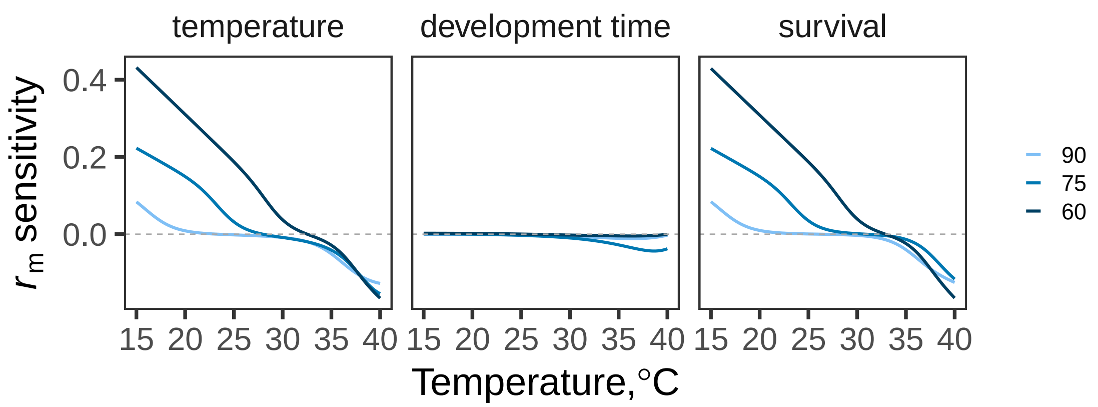

**Figure S9: Seasonal mean temperatures and relative humidity across Africa and South Asia under historical climate conditions (1970–2000).** Panels (a) and (c) show mean annual temperatures by season across Africa and India for 1970–2000, respectively. Panels (b) and (d) show mean annual relative humidity by season across Africa and India for the same period, respectively.

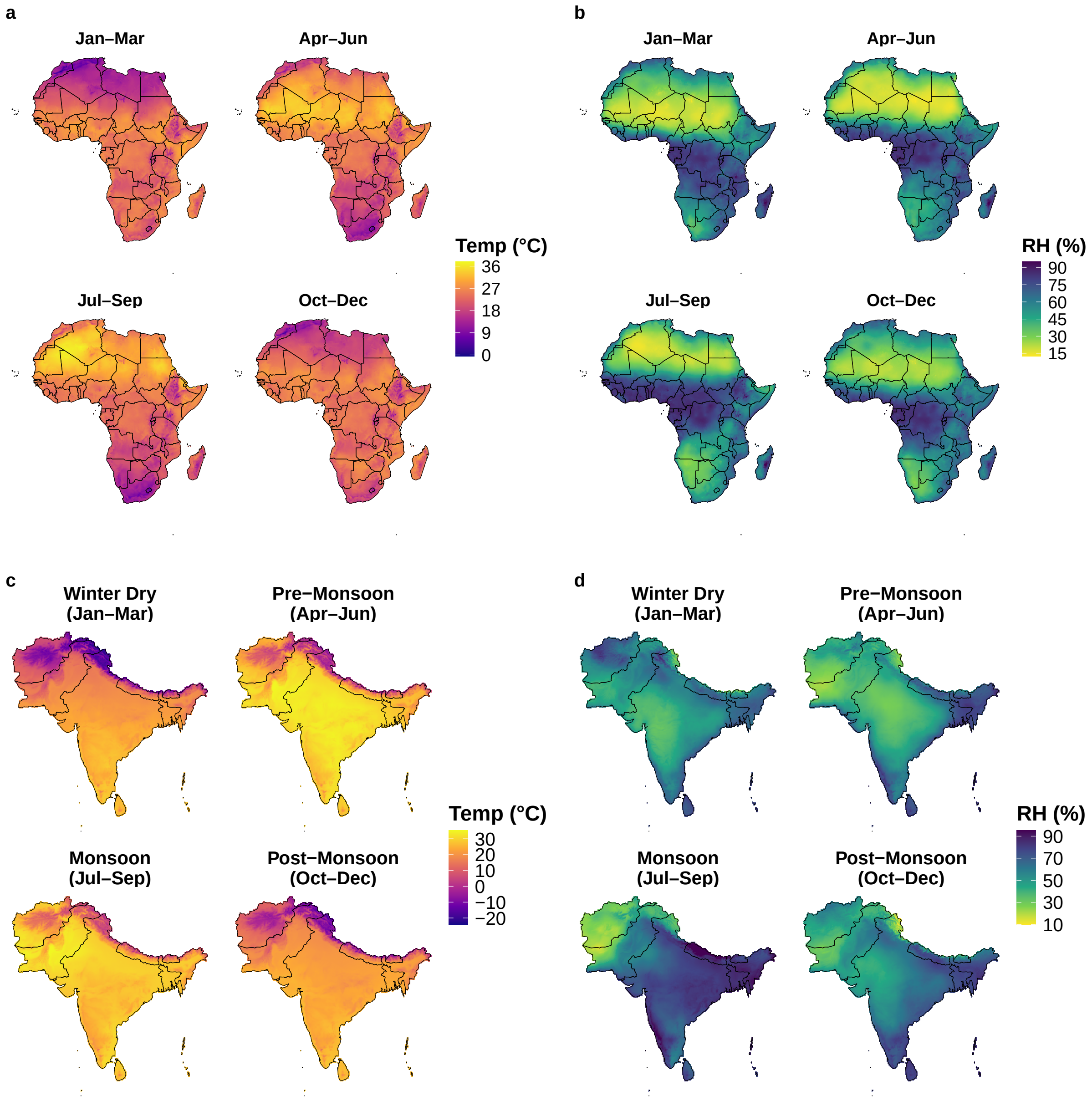
